# AI-ASSISTED NANOMEDICINE REVEALS SYSTEMIC IMMUNE IMPACT IN OVARIAN CANCER METASTASIS

**DOI:** 10.64898/2026.10.12.758609

**Authors:** Riya Manandhar, Haya Tchalabi, Yasodha Kesavan, Sam Koroma, Laura Holman, Rupa Haldavnekar

**Affiliations:** Data Science and Artificial Intelligence, Gallogly College of Engineering, The University of Oklahoma - 202 W Boyd St., Rm 409, Norman, Oklahoma; School of Electrical and Computer Engineering, Gallogly College of Engineering, The University of Oklahoma-Devon Energy Hall, 150 110 W. Boyd St., Norman, Oklahoma; OU Health Stephenson Cancer Center, The University of Oklahoma, Harold Hamm Health Campus

## Abstract

Ovarian cancer, a gynecological malignancy, poses a significant challenge as researchers struggle to comprehend how tumor cells communicate with the immune system. Ovarian tumors contain both bulk cancer cells and cancer stem cells (CSCs), which are linked to chemoresistance, relapse, and immune escape. However, it remains unclear whether these two populations affect immune cells differently. To investigate this, a study used surface-enhanced Raman spectroscopy (SERS) combined with machine learning to examine the biochemical changes in peripheral blood mononuclear cells (PBMCs) after co-culture with OVCAR4 ovarian cancer cells or OVCAR4-derived CSC-enriched populations. PBMCs and tumor cells were co-cultured for 48 hours in an indirect Transwell system, allowing exchange of soluble factors without direct cell contact. SERS spectra were acquired on gold-coated nickel substrates and analyzed using principal component analysis, t-test, and a linear support vector machine (SVM). The results showed that PBMCs co-cultured with CSCs and PBMCs co-cultured with OVCAR4 cells exhibited distinct spectral profiles. Moreover, the SVM demonstrated a high cross-validated accuracy of 98.9% in separating these two populations. PBMCs exposed to CSCs exhibited higher adenine and carbohydrate signals, while those exposed to OVCAR4 cells displayed higher protein and lipid signals. Interestingly, the tumor cells themselves exhibited the same pattern at baseline. Moreover, six out of the ten most discriminating bands were shared between the tumor and PBMC comparisons, indicating a consistent direction of change. These findings suggest that tumor-derived soluble factors leave a distinct biochemical signature on PBMCs. Label-free SERS combined with machine learning offers a sensitive approach for studying tumor-immune communication and developing blood-based tools to monitor ovarian cancer.

## Introduction

Ovarian cancer remains one of the major causes of gynecological cancer-related deaths. The high mortality is associated with late diagnosis, recurrence, treatment resistance, and the heterogeneous nature of the disease^1,2^. Although immunotherapy has improved treatment outcomes in several cancers, its effectiveness in ovarian cancer has been limited. One reason is the complex interaction between ovarian cancer cells and the immune system, which can lead to suppression or alteration of antitumor immune responses^3,4^. Understanding how ovarian cancer cells interact with immune cells is therefore important for identifying biological changes associated with disease progression and treatment response^4–6^.

Ovarian cancer is highly heterogeneous and contains multiple cancer-cell populations with distinct molecular and functional properties^5,6^. Cancer stem-like cells (CSCs) are a subpopulation with properties such as self-renewal, tumor initiation, treatment resistance, and the ability to contribute to tumor recurrence^7,8^. Importantly, ovarian CSCs can interact with immune cells and alter immune surveillance and immune-cell function. This suggests that the immune response to a CSC-enriched population may be different from the response to the parental cancer-cell population^7,8^. However, the biochemical changes that occur in peripheral immune cells following interaction with these different ovarian cancer cell populations are not fully understood^4,9^.

Peripheral blood mononuclear cells (PBMCs) include circulating lymphocytes and monocytes and represent an accessible source of peripheral immune cells^4^. Cancer-associated signals can alter the phenotype and function of immune cells, and these changes may provide information about the interaction between the tumor and the peripheral immune system^4,10^. Therefore, this study investigates whether interactions with different ovarian cancer cell populations produce measurable biochemical changes in PBMCs and whether PBMCs exposed to parental OVCAR4 ovarian cancer cells exhibit biochemical profiles distinct from those exposed to OVCAR4-derived CSC-enriched cells^9,11^.

This concept is also relevant to our longer-term goal of developing a liquid-biopsy approach for ovarian cancer. Liquid biopsy provides a minimally invasive way to obtain tumor-associated information from blood and other body fluids and has been investigated for cancer detection, disease monitoring, and treatment-response assessment. Most liquid-biopsy approaches focus on circulating tumor cells, extracellular vesicles, circulating nucleic acids, proteins, or other tumor-associated molecules. However, circulating immune cells are also continuously exposed to tumor-associated signals^12,13^. We propose that biochemical changes occurring in PBMCs following tumor interaction may contain additional information about the biological state of the tumor. Before this possibility can be evaluated in patients, it is important to determine in a controlled experimental system whether different ovarian cancer cell populations can produce distinguishable biochemical changes in PBMCs.

Surface-enhanced Raman spectroscopy (SERS) provides a sensitive method for studying these biochemical changes without requiring a predefined panel of molecular targets. SERS enhances Raman scattering through interactions between molecules and plasmonic metallic nanostructures and generates spectral fingerprints containing information associated with proteins, lipids, nucleic acids, carbohydrates, and other cellular components^12–14^. SERS has increasingly been investigated for cancer liquid-biopsy applications using blood and other biofluids, circulating tumor cells, extracellular vesicles, nucleic acids, and proteins. Because the resulting spectra contain many overlapping biochemical features, multivariate analysis and machine-learning methods can be used to identify spectral patterns and determine whether different biological groups can be distinguished^12,15^.

In the present study, we used parental OVCAR4 ovarian cancer cells and OVCAR4-derived CSC-enriched cells to investigate how different ovarian cancer cell populations influence the biochemical profile of PBMCs. PBMCs were exposed to either parental OVCAR4 cells or CSC-enriched cells, and SERS spectra were collected from the PBMCs following these interactions. The resulting spectral profiles were analyzed using multivariate and machine-learning approaches to determine whether exposure to the two cancer cell populations produced distinguishable biochemical changes in PBMCs. We hypothesized that parental OVCAR4 cells and CSC-enriched cells would influence PBMCs differently, resulting in distinct SERS spectral signatures. This controlled cell-based study provides an initial step toward identifying tumor-associated biochemical changes in peripheral immune cells that can subsequently be evaluated in patient-derived blood samples for potential SERS-based ovarian cancer liquid-biopsy applications.

## Materials and Methods

### Cell culture and PBMC indirect co-culture

Human OVCAR4 ovarian cancer cells were maintained in RPMI 1640 supplemented with 10% fetal bovine serum (FBS) at 37°C in a humidified atmosphere containing 5% CO_2_.

OVCAR4-derived cancer stem cell (CSC)-enriched populations were generated under serum-free sphere-forming conditions using DMEM/F12 supplemented with 1× B27 (Thermo Fisher Scientific, Cat. No. 17504044), recombinant human epidermal growth factor (EGF; 20 ng/mL), recombinant human basic fibroblast growth factor (bFGF/FGF2; 20 ng/mL), and heparin (2 µg/mL; STEMCELL Technologies, Cat. No. 07980). Cells were seeded at 10,000 cells per well in ultra-low-attachment 96-well plates and maintained at 37°C in a humidified atmosphere containing 5% CO_2_. One spheroid was generated per well. Spheroids from three wells were carefully collected and transferred to a standard 24-well culture plate for subsequent indirect co-culture experiments.

Human peripheral blood mononuclear cells (PBMCs) were obtained from STEMCELL Technologies (Cat. No. 70025.1). An indirect co-culture system was established using 0.4-µm-pore Transwell inserts, which physically separated the PBMCs from the tumor cells while permitting exchange of soluble factors and extracellular vesicles. PBMCs and OVCAR4 parental cells or CSC-enriched populations were co-cultured at a 1:1 ratio for 48 h. OVCAR4 cells/CSC-enriched populations were maintained in the lower compartment, while PBMCs were placed in the upper Transwell insert.

Experimental groups consisted of OVCAR4 control, OVCAR4 + PBMC, CSC control, CSC + PBMC, PBMC control, PBMC + OVCAR4, and PBMC + CSC. Following 48 h of co-culture, tumor cells/CSC-enriched populations and PBMCs were collected separately, washed, and prepared for Raman spectroscopic analysis.

### SERS substrate

Gold-coated nickel substrates were fabricated using Glancing Angle Deposition (GLAD) in a Kurt J. Lesker thermal evaporator. Unpolished nickel sheets were cleaned by sonication using Acetone, IPA, and DI water for four minutes each, followed by a 5-minute oxygen plasma treatment under vacuum to remove any remaining contaminants using a Technics Planar Etch II Plasma Asher.

Custom inclined platforms were used to position the samples at an 85° angle relative to the substrate norm; Those were fabricated from the same nickel material as the substrate to minimize the introduction of additional materials that could contribute to outgassing or contamination during thermal deposition. The substrates were rotated during deposition to promote a more uniform distribution of the nanoparticles.

Gold was deposited at a thickness of 20 nm. The resulting AU/Ni substrates were then used for Surface Enhanced Raman Spectroscopy measurements.

### Data Collection

Raman spectra were acquired using a Qontor InVia Raman confocal micro-spectroscope (Renishaw). To maintain consistency and comparability across all samples and time points, all instrumental parameters were kept constant throughout the study, including the laser excitation wavelength (785nm), laser power at the sample, integration time, and objective magnification. Spectral acquisition was performed in the Raman fingerprint region (approximately 600–1800 cm^−1^), which encompasses the biomolecular vibrational signatures most informative for characterizing proteins, lipids, nucleic acids, and metabolites^14^. Approximately (492-1072) spectra were collected per sample to adequately capture the intra-sample biochemical variability. Before further analysis, visual inspection was used to identify and exclude spectra exhibiting acquisition-related artifacts, such as cosmic ray spikes, to ensure the downstream data quality and reliability.

### Spectral Preprocessing

SERS spectra were acquired and saved as .spc files. Baseline correction and Savitzky–Golay smoothing were applied in Spectragryph software, and the processed spectra were exported as CSV files (1759.8–660.6 cm^−1^, 1015 points per spectrum). In Python, exported baseline curves and all spectra were mapped onto a common Raman-shift axis to correct small (∼0.001 cm^−1^) rounding differences between files. Cosmic-ray spikes were removed by comparing each spectrum with its five-point moving median and replacing any point that deviated by more than five standard deviations of that spectrum’s deviations with the median value. This altered only individual points, and no spectra were discarded. Spectra were then vector-normalized by dividing each by its Euclidean (L2) norm. This corrects for overall intensity differences between spectra without assuming that any single band is constant across cell types. Because the groups differed in size (494 to 1072 spectra), each comparison was class-balanced by random subsampling without replacement (fixed random seed) to the size of the smallest group, capped at 500. This gave 494 spectra per group in comparisons 1 and 2 and 500 per group in comparisons 3 and 4.

### Data Analysis

All analysis were conducted in Python 3.13 on the pre-processed spectra. Principal component analysis (PCA) was employed to visualize spectral variation and differentiate between sample groups. Score plots were generated for the component pair exhibiting the most distinct separation. To pinpoint the Raman bands responsible for this separation, the local maxima and minima of the PC1 loading curve were identified using scipy.signal.find_peaks. Subsequently, the normalized intensity at each of these bands was compared across groups using Welch’s t-test. The direction and size of change at each band were expressed as the percent change in mean normalized intensity. Bands were assigned to biomolecule classes (protein, lipid, nucleic acid, carbohydrate) using published Raman band assignments^14^, and the percent changes were averaged within each class to summarise class-level trends. To test whether the groups could be distinguished from the full spectrum, a linear-kernel support vector machine (SVM) was trained on all normalized intensities and evaluated by stratified five-fold cross-validation, and the SVM coefficients were plotted against Raman shift to show which bands the classifier relied on. Analysis and figures used pandas, NumPy, SciPy, scikit-learn, and Matplotlib.

## Result and Discussion

Prior to applying the fabricated substrates to biological measurements, crystal violet (CV) with 100 µM was used as a reporter to evaluate the rationale for both SERS-based detection and the Glancing Angle Deposition (GLAD) fabrication approach.

Raman spectra were collected from CV on three surfaces: 20 nm GLAD Au/Ni substrate, and a 20 nm normal-incidence Au/Ni substrate, both for SERS measurements, and bare silicon for Raman with no enhancement. All three cases underwent the same acquisition conditions of a 100% laser power, 1 second acquisition time and 1 accumulation.

As shown in Fig 1A, the GLAD substrate produced substantially stronger CV Raman peaks for crystal violet than the normal-incidence Au substrate, with bare silicon substrate showing the dimmest signal. The weak response observed on bare silicon illustrates the limited detectability of dilute CV under the same acquisition conditions without a SERS-active surface.

**Figure 1.**
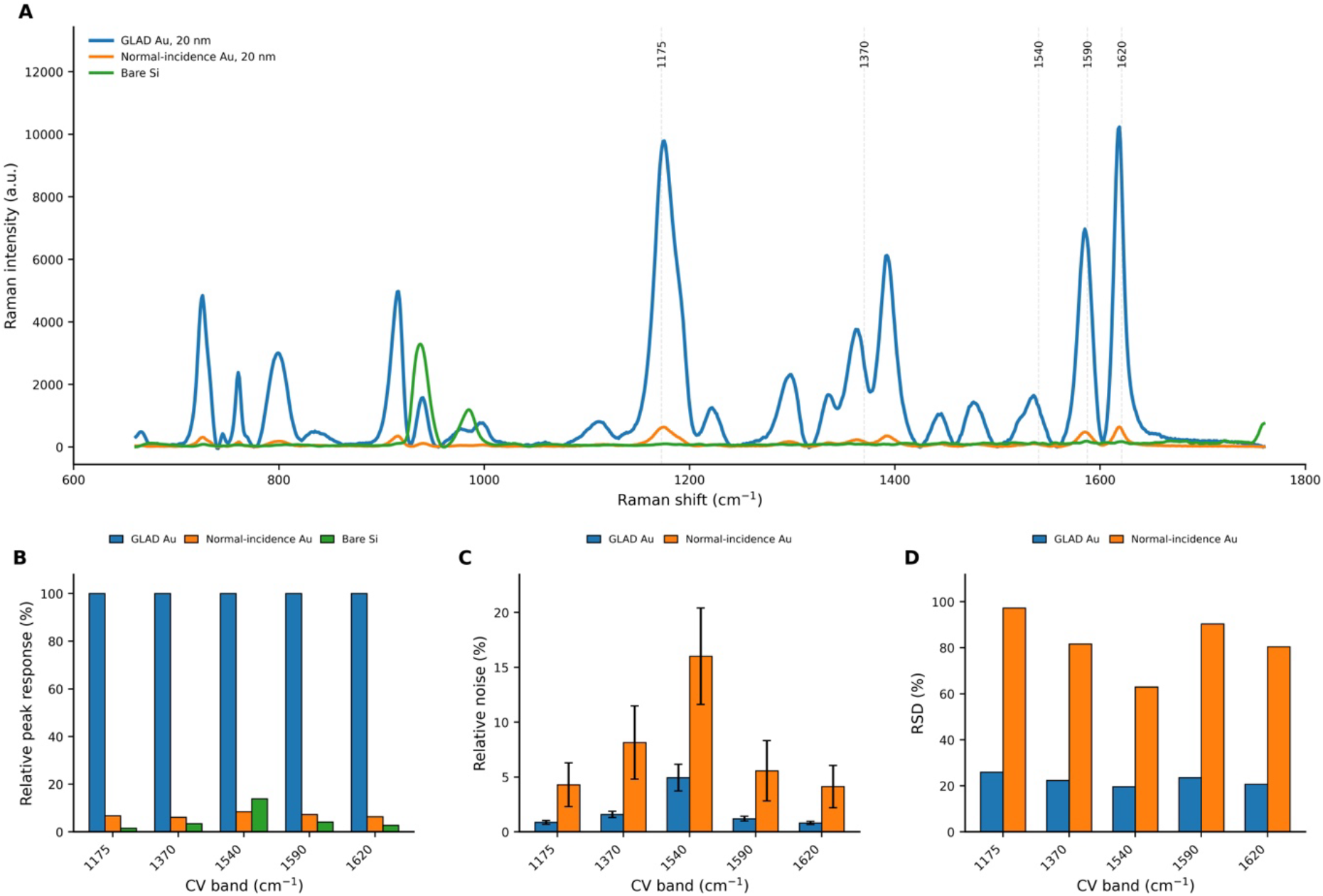
Preliminary SERS evaluation of 20 nm GLAD Au/Ni substrates using crystal violet. (A) Mean baseline-corrected Raman spectra of CV measured on 20 nm GLAD Au/Ni, 20 nm normal-incidence Au/Ni, and bare silicon substrates. (B) Relative peak response at selected CV bands, expressed as a percentage of the corresponding GLAD peak intensity. (C) Relative spectral noise for the GLAD and normal-incidence Au substrates. For each spectrum, the standard deviation of Raman intensity within a peak-free region (1700–1780 cm^−1^) was used as an estimate of spectral noise. This value was divided by the intensity of each selected CV band and expressed as a percentage, such that lower values indicate a smaller noise contribution relative to the Raman peak intensity. (D) Relative standard deviation (RSD) of selected CV peak intensities across replicate measurements, calculated for each peak as RSD (%) = SD_peak / mean peak intensity × 100, used to assess measurement variability. The GLAD substrate exhibited both lower relative noise and lower RSD than the normal-incidence Au substrate.

Bare Silicon provides a reference point for evaluating the CV signal without gold deposition. Since the signal intensity is higher on the gold-deposited substrates, especially for GLAD Au, these differences are consistent with the behavior contributed by the gold-associated architecture. Compared with metallic nanostructures, flat or bare silicon relies heavily on spontaneous Raman scattering and is very ineffective at capturing molecular changes, as the substrate cannot manipulate incoming light. Furthermore, for our experiment, the cells will adhere to the flat surface of bare silicon, which is why our figure shows a relatively low signal-to-noise ratio for this substrate. We would need longer acquisition times for the Silicon peak to dominate, which can cause prothermal damage to cells^17^.

Furthermore, quantification of representative CV bands showed that the signal measured on the normal-incidence Au substrate corresponded to only approximately 6–8% of the GLAD response for most selected bands, while the bare silicon response was generally only a few percent of the GLAD signal (Fig 1B). These results support the use of a nanostructured Au surface for enhancing weak Raman signals and indicate an additional benefit from the oblique-angle GLAD geometry beyond Au deposition alone. The array of nanocolumns on GLAD Au provides us with a dense plasmonic structure that interacts directly with the cellular envelope and enables us to capture non-discriminatory, label-free biomolecular changes in ovarian cancer populations and PBMCs.

Figure 1 therefore highlights the variations between the three substrates. These differences between GLAD Au and Normal-incidence Au may be attributed to how different their surface morphologies are and the spatial distribution of electromagnetic field-enhancing regions. Signal quality and measurement variability were also assessed for the two Au-coated substrates. Relative noise was consistently lower for the GLAD substrate across the selected CV bands, ranging from approximately 0.8–4.9%, compared with approximately 4.1–16.0% for the normal-incidence Au substrate (Fig 1C). In addition, the GLAD substrate showed substantially lower relative standard deviation (RSD), approximately 20–26%, compared with approximately 63–97% for the normal-incidence Au substrate (Fig 1D). Together, these preliminary measurements indicate that the GLAD substrate provides not only greater Raman signal intensity but also improved signal quality and measurement consistency compared with a conventional 20 nm Au coating

For our experiment, the quality of the signals from the substrate and measurement variability are important. When we interpret the spectra from the ovarian cancer populations and their influence on PBMCS, variations that stem from the substrates could cause signal intensity that overlap with observations of the differences between the biological groups.

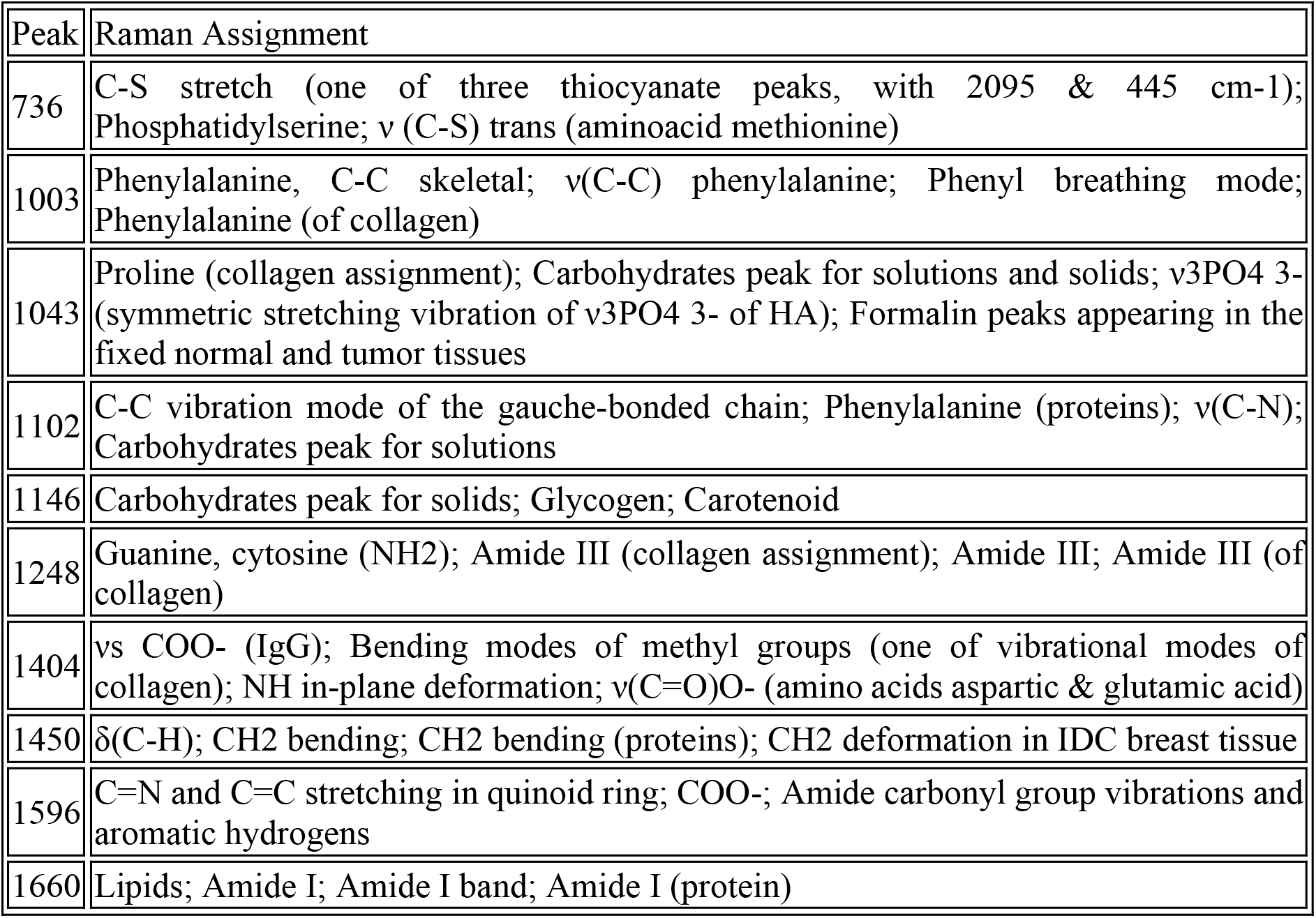

SERS separated the two cancer cell populations clearly. The mean spectra (Fig. 2A) differ in shape, and PCA places OVCAR4 and CSC spectra in largely distinct regions of the PC1–PC3 plane, with PC1 alone accounting for 40.8% of the variance (Fig. 2B). The ten most discriminating bands (Fig. 2C) all differed significantly between the groups (p < 10^−27^ for each), so the separation rests on many bands rather than one. Spectra were vector-normalized, so differences reflect relative spectral contributions rather than absolute amounts.

**Figure 2.**
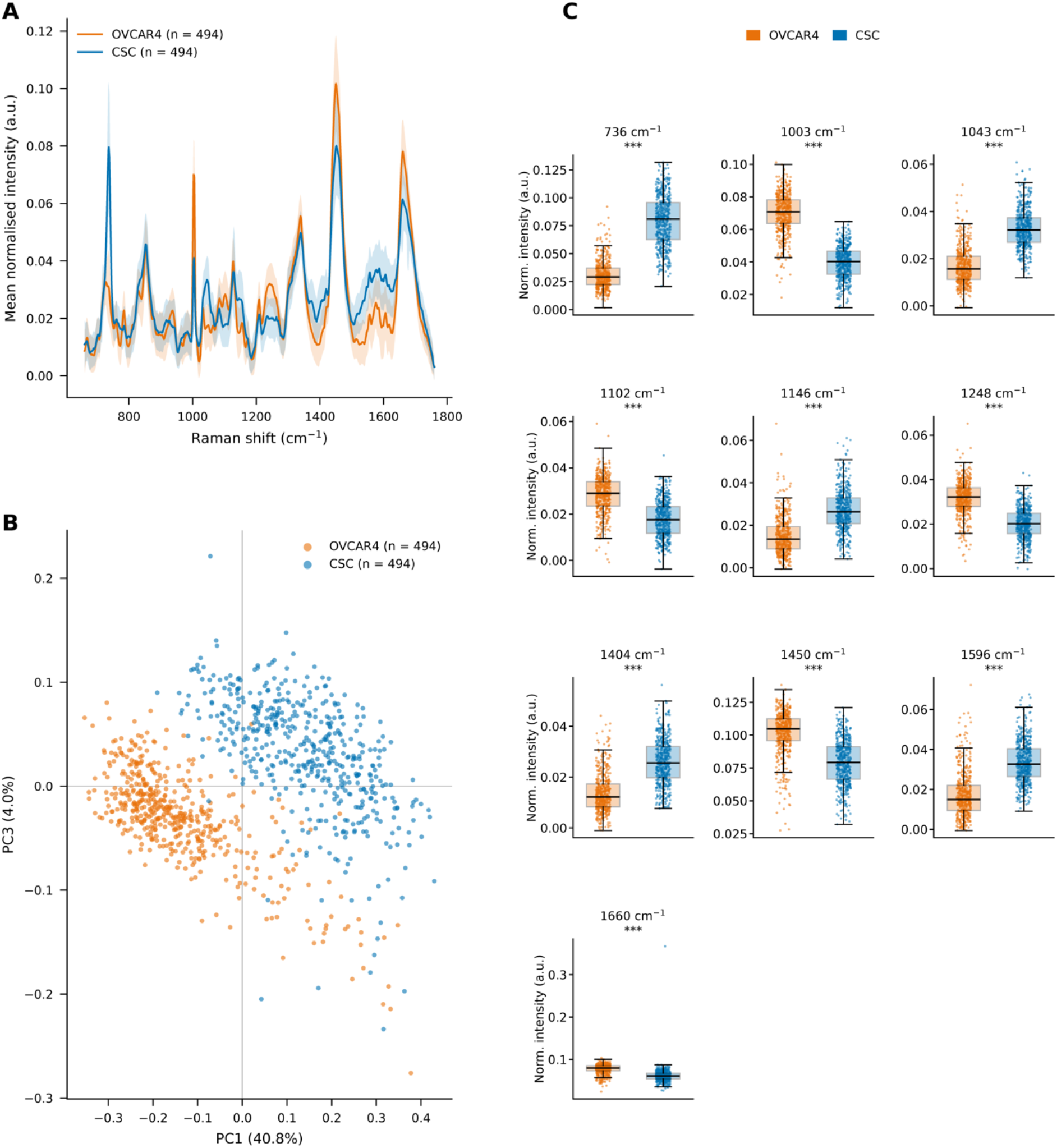
SERS discriminates OVCAR4 cells from OVCAR4-derived cancer stem cells (CSCs). (A) Mean spectra of OVCAR4 and CSCs. (B) Principal component analysis (PCA). (C) t-test of the ten most significant bands.

The two populations exhibited distinct biomolecular classes dominating their spectra. OVCAR4 contributed significantly at 1003 cm^−1^ (phenylalanine ring breathing), 1248 cm^−1^ (amide III), 1450 cm^−1^ (CH_2_/CH_3_ deformation of proteins and lipids), and 1660 cm^−1^ (amide I, which also encompasses C=C stretching of unsaturated lipids), resulting in a spectrum predominantly characterized by protein-associated bands. In contrast, CSC contributed more at 736 cm^−1^, the most discriminating band (t ≈ 42), a region assigned to adenine ring breathing and phospholipid head-group vibrations (choline, phosphatidylserine). Additionally, CSC was present at 1043 and 1146 cm^−1^, which fall within regions associated with carbohydrates, glycogen, proline, and C–C/C–N vibrations^14^. Furthermore, CSC was higher at 1404 and 1596 cm^−1^ (COO^−^/CH_3_ bending and C=C/aromatic ring modes). Given the overlap of several of these bands across multiple biomolecule classes, we interpret them as shifts in overall composition rather than definitive evidence for a single molecule.

This pattern is consistent with reports that cancer stem-like cells differ metabolically from the bulk tumor population. In ovarian cancer, CSCs exhibit increased lipid desaturation, a finding confirmed by Raman-based imaging and mass spectrometry. This lipid desaturation is functionally essential for the stemness of CSCs^7^. CSCs also show metabolic plasticity, switching between glycolysis and oxidative phosphorylation and drawing on fatty-acid synthesis and uptake^18^, and breast CSC models activate glycogen breakdown and glycolysis ^19^. The higher CSC contribution in lipid/phospholipid-, nucleic-acid- and carbohydrate/glycogen-associated regions in our data fits this general picture, although we did not measure these pathways directly, and ref. ^19^ was obtained in breast rather than ovarian models. We note that our comparison is between two cell populations measured on the same substrate, so it does not by itself identify the molecules responsible.

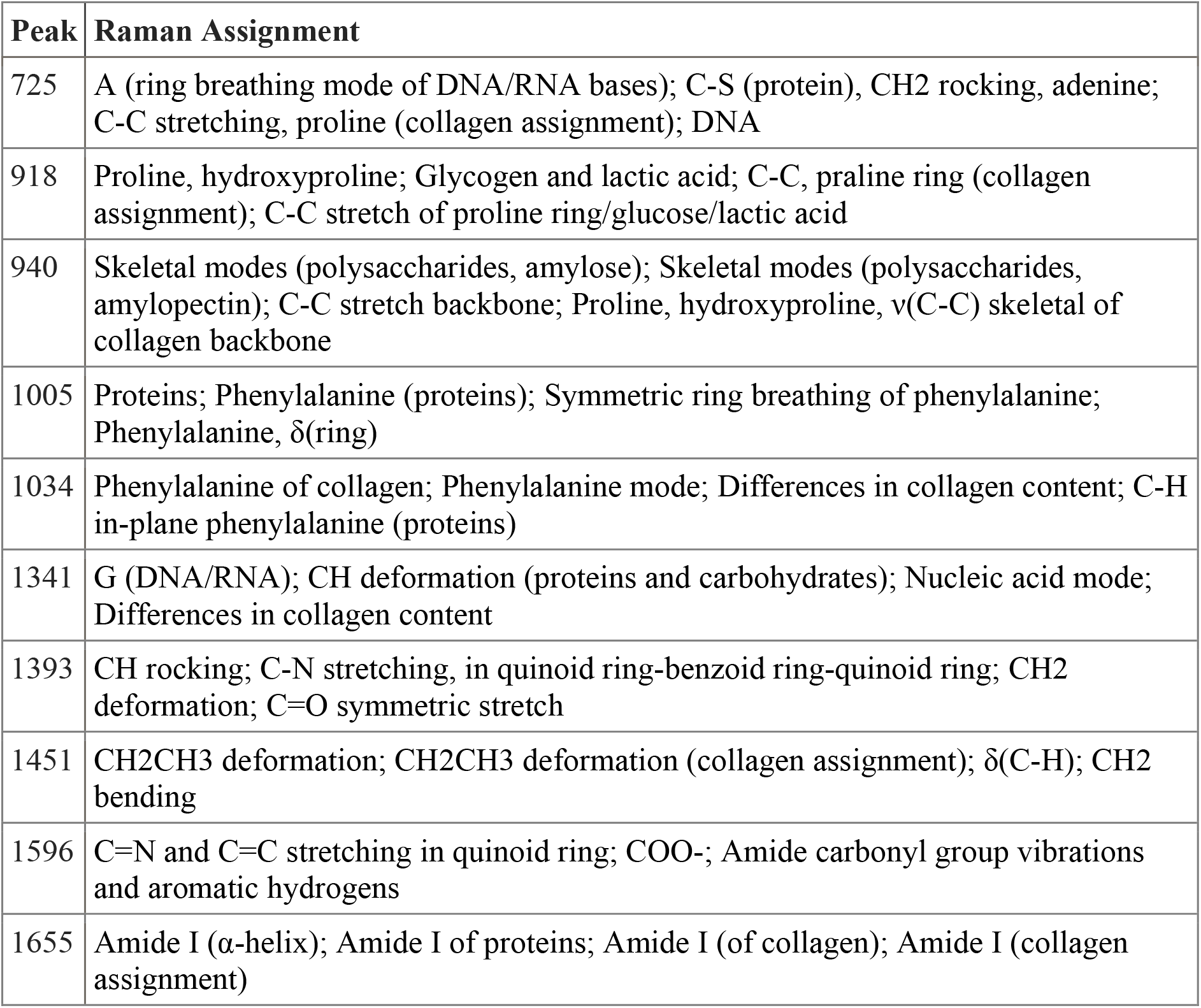

| Peak | Raman Assignment |
| --- | --- |
| 725 | A (ring breathing mode of DNA/RNA bases); C-S (protein), CH<sub>2</sub> rocking, adenine; C-C stretching, proline (collagen assignment); DNA |
| 918 | Proline, hydroxyproline; Glycogen and lactic acid; C-C, praline ring (collagen assignment); C-C stretch of proline ring/glucose/lactic acid |
| 940 | Skeletal modes (polysaccharides, amylose); Skeletal modes (polysaccharides, amylopectin); C-C stretch backbone; Proline, hydroxyproline, $\nu$(C-C) skeletal of collagen backbone |
| 1005 | Proteins; Phenylalanine (proteins); Symmetric ring breathing of phenylalanine; Phenylalanine, $\delta$(ring) |
| 1034 | Phenylalanine of collagen; Phenylalanine mode; Differences in collagen content; C-H in-plane phenylalanine (proteins) |
| 1341 | G (DNA/RNA); CH deformation (proteins and carbohydrates); Nucleic acid mode; Differences in collagen content |
| 1393 | CH rocking; C-N stretching, in quinoid ring-benzoid ring-quinoid ring; CH<sub>2</sub> deformation; C=O symmetric stretch |
| 1451 | CH<sub>2</sub>CH<sub>3</sub> deformation; CH<sub>2</sub>CH<sub>3</sub> deformation (collagen assignment); $\delta$(C-H); CH<sub>2</sub> bending |
| 1596 | C=N and C=C stretching in quinoid ring; COO<sup>-</sup>; Amide carbonyl group vibrations and aromatic hydrogens |
| 1655 | Amide I (α-helix); Amide I of proteins; Amide I (of collagen); Amide I (collagen assignment) |

SERS distinguished PBMCs before and after co-culture with OVCAR4. PCA placed untreated and co-cultured PBMC spectra in largely distinct regions of the PC1–PC3 plane, with PC1 accounting for 35.9% and PC3 for 4.3% of the variance (Fig. 3A); a subset of co-cultured spectra extended towards the untreated cluster, so separation was not complete in this projection. The ten most discriminating bands (Fig. 3B) all differed significantly between the groups (p < 10^−39^ for each), so the separation rests on many bands rather than one. Spectra were vector-normalized, so differences are described as relative spectral contributions rather than absolute amounts.

**Figure 3.**
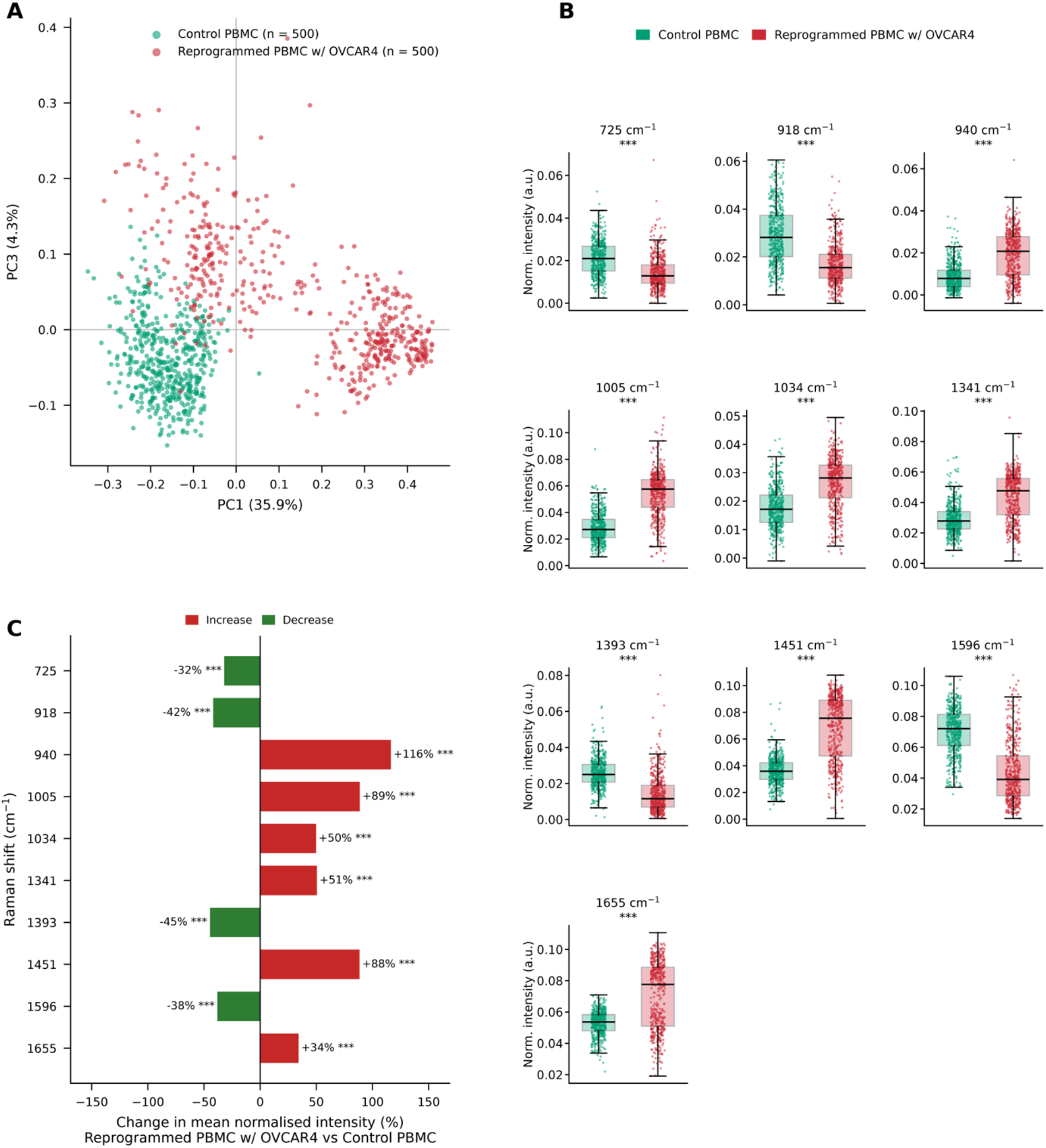
Control PBMCs and reprogrammed PBMC with OVCAR4. (A) Principal component analysis (PCA). (B) t-test of the ten most significant bands. (C) Direction of change (% change in co-cultured vs control PBMCs).

Co-cultured PBMCs contributed more at 1005 cm^−1^ (phenylalanine ring breathing, +89%), 1034 cm^−1^ (phenylalanine C–H bending, +50%), 1655 cm^−1^ (amide I, +34%), 1451 cm^−1^ (CH_2_/CH_3_ deformation of proteins and lipids, +88%), 940 cm^−1^ (C–C backbone stretch of proline and valine, which also falls in a glycogen-associated region, +116%) and 1341 cm^−1^ (guanine and CH deformation, +51%) (Fig. 3C). In contrast, untreated PBMCs contributed more at 725 cm^−1^ (adenine ring breathing, C–S/CH_2_ rocking, −32% after co-culture), 918 cm^−1^ (proline, hydroxyproline and glycogen-associated modes, −42%), 1393 cm^−1^ (non-specific CH rocking, −45%) and 1596 cm^−1^ (C=C and aromatic ring modes, −38%)^14^. The increases were therefore concentrated in protein-assigned bands, while the decreases fell in bands of mixed or less specific assignment. Given the overlap of several of these bands across biomolecule classes, we interpret them as shifts in overall composition rather than as evidence for a single molecule.

This protein-weighted pattern is consistent with a change in immune-cell state, because Raman spectra of immune cells change with state: naïve and activated T cells can be classified from their spectra^15^, macrophage activation can be detected from Raman spectra combined with morphology^20^, and activation-induced changes in CD8? T cells include increased protein bands at 1002 and 1234 cm^−121^. The spectra do not identify which state or which cell type changed, and because PBMCs are a mixed population, shifts in the mean spectrum may also reflect changes in cell composition. The data do not show the lipid-loading signature that has been linked to immune dysfunction in tumor settings^22,23^, as the cholesterol/lipid-associated 725 cm^−1^ band decreased and nucleic-acid-associated bands moved in opposite directions (725 down, 1341 up). The bands that increased include those that were stronger in OVCAR4 itself (1003–1005, 1450 and 1660 cm^−1^; Fig. 3), which is consistent with uptake of cancer-derived material, since ovarian cancer exosomes are known to alter lymphocyte signaling^21^, but is equally consistent with residual cancer cells or debris in the PBMC preparation, which spectra alone cannot separate. We note that these changes were measured on one substrate after co-culture, so they do not by themselves identify the molecules or mechanism responsible.

| Peak | Raman Assignment |
| --- | --- |
| 728 | Adenine ring breathing (DNA/RNA bases); C–S (protein), CH<sub>2</sub> rocking |
| 996 | C–O stretch of ribose; C–C stretch |
| 1004 | Phenylalanine ring breathing |
| 1013 | Carbohydrates; phenylalanine |
| 1045 | Carbohydrates; proline; glycogen |
| 1310 | CH<sub>3</sub>/CH<sub>2</sub> twisting and wagging (lipids, proteins) |
| 1453 | CH<sub>2</sub>/CH<sub>3</sub> deformation (proteins, lipids) |
| 1600 | C=C in-plane bending of phenylalanine and tyrosine |

PBMCs were co-cultured with OVCAR4-derived CSCs, and their spectra were analyzed using PCA. The PCA placed untreated and co-cultured PBMC spectra in distinct regions of the PC1– PC2 plane, with PC1 accounting for 29.2% and PC2 for 14.0% of the variance (Fig. 4A). Only ten bands were detected in the PC1 loading, as shown in Fig. 4B. Eight of these ten bands differed significantly between the groups (p = 2.6 × 10^−173^ to 7.0 × 10^−3^), while the bands at 789 and 855 cm^−1^ did not (p = 0.20 and 0.27). Therefore, the separation between the groups is primarily based on a smaller set of bands compared to the OVCAR4 condition.

**Figure 4.**
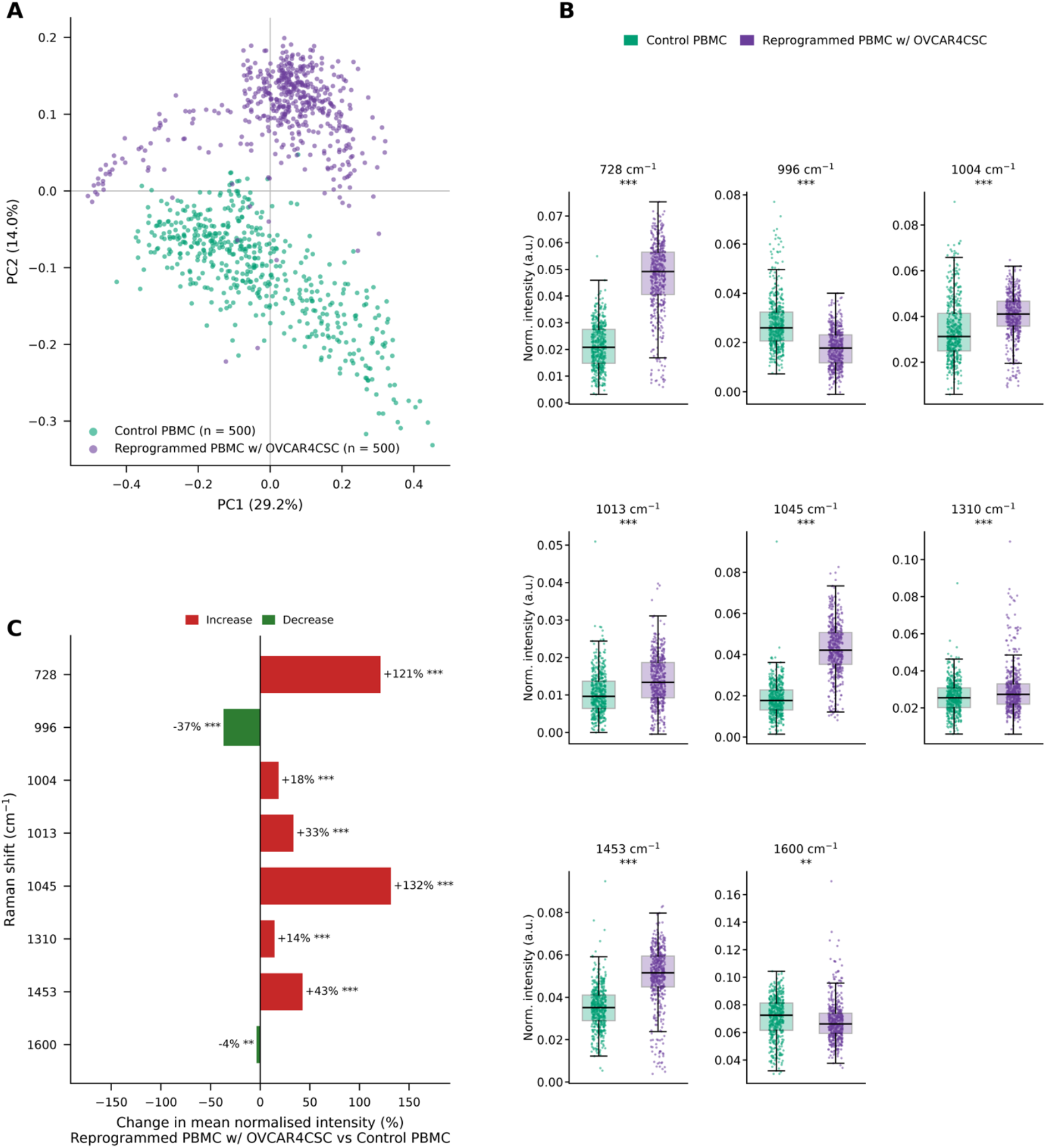
Control PBMCs and reprogrammed PBMCs with OVCAR4-derived CSCs. (A) Principal component analysis (PCA). (B) t-test of the significant bands. (C) Direction of change (% change in reprogrammed PBMCs vs control PBMCs).

Co-cultured PBMCs contributed more at 1045 cm^−1^ (a carbohydrate/glycogen-associated region that also includes proline modes, +132%) and 728 cm^−1^ (adenine ring breathing and phospholipid-associated modes, +121%), followed by 1453 cm^−1^ (CH_2_ deformation of proteins and lipids, +43%), 1013 cm^−1^ (phenylalanine-associated, +33%), 1004 cm^−1^ (phenylalanine ring breathing, +18%) and 1310 cm^−1^ (CH_3_/CH_2_ twisting of lipids and collagen, +14%) (Fig. 4C). In contrast, untreated PBMCs contributed more at 996 cm^−1^ (C–O ribose/C–C stretch; phenylalanine-associated region, −37% after co-culture), and the 1600 cm^−1^ aromatic C=C band changed only slightly (−3.6%)^14,24^. Compared with the OVCAR4 condition, the increases at the phenylalanine (1004 cm^−1^, +18% vs +89%) and CH_2_/CH_3_ (1453 cm^−1^, +43% vs +88%) bands were smaller, whereas the largest increases here were at 728 and 1045 cm^−1^. Given the overlap of several of these bands across biomolecule classes, we interpret them as shifts in overall composition rather than as evidence for a single molecule.

The two largest increases fall at the bands that were stronger in CSCs themselves than in OVCAR4 (736 and 1043 cm^−1^; Fig. 2), which is consistent with PBMCs taking up cancer-stem-cell-derived material, as ovarian cancer exosomes are known to alter lymphocyte signaling^10^. It is equally consistent with residual cancer cells or debris in the PBMC preparation, which spectra alone cannot separate. Ovarian CSCs are reported to carry increased lipid desaturation^7^, and the rise of the lipid/phospholipid-associated 728 and 1310 cm^−1^ bands is of the type that has been linked to lipid accumulation and dysfunction in tumor-exposed immune cells^22,23^, but a single Raman band cannot show lipid accumulation, and the same 725 cm^−1^ band fell in the OVCAR4 condition (Fig. 3). More generally, Raman spectra of immune cells change with cell state^15,20,21^, so a spectral shift is consistent with a change in state without identifying which state or which cell type changed, and because PBMCs are a mixed population, part of the shift may reflect changes in cell composition. We note that these changes were measured on one substrate after co-culture, so they do not by themselves identify the molecules or mechanism responsible.

| Peak | Raman Assignment |
| --- | --- |
| 726 | C-S (protein), CH<sub>2</sub> rocking, adenine; A (ring breathing mode of DNA/RNA bases); C-C stretching, proline (collagen assignment); A |
| 938 | C-C stretch backbone; Proline, hydroxyproline, $\nu$(C-C) skeletal of collagen backbone; Glycogen; Proline (collagen type I) |
| 1005 | Proteins; Phenylalanine (proteins); Symmetric ring breathing of phenylalanine; Phenylalanine, $\delta$(ring) |
| 1014 | Carbohydrates peak for solids; Stretching C-O ribose; Amphetamine has a group of three bands at this region |
| 1042 | Proline (collagen assignment); Carbohydrates peak for solutions and solids; Formalin peaks appearing in the fixed normal and tumor tissues; ν3PO<sub>4</sub> 3-(symmetric stretching vibration of ν3PO<sub>4</sub> 3- of HA) |
| 1081 | ν1CO<sub>3</sub> 2-, ν3PO<sub>4</sub> 3-, $\nu$(C-C) skeletal of acyl backbone in lipid (gauche conformation); Nucleic acids; Carbohydrates peak for solutions; Carbohydrate residues of collagen |
| 1341 | G (DNA/RNA); CH deformation (proteins and carbohydrates); Nucleic acid mode; Differences in collagen content |
| 1450 | $\delta$(C-H); CH<sub>2</sub> bending; CH<sub>2</sub> bending (proteins); CH<sub>2</sub> deformation in IDC breast tissue |
| 1597 | C=N and C=C stretching in quinoid ring; Phenylalanine; $\delta$(C=C), phenylalanine (protein assignment); COO- |
| 1655 | Amide I (α-helix); Amide I of proteins; Amide I (of collagen); Amide I (collagen assignment) |

PBMCs co-cultured with OVCAR4 and those co-cultured with OVCAR4-derived CSCs showed distinct SERS signatures. In the PC1–PC3 plane, the CSC-exposed PBMCs formed a compact cluster, whereas the OVCAR4-exposed PBMCs spread over two sub-populations along PC1 (PC1 30.9%, PC3 7.3%; Fig. 5A). Of the 20 bands detected in the PC1 loading, the ten most discriminating all differed significantly between the groups (p = 2.3 × 10^−233^ to 2.6 × 10^−24^; Fig. 5B). A linear support vector machine trained on the full spectrum classified the two groups with 98.9 ± 0.4% cross-validated accuracy, with 492 of 500 OVCAR4-exposed and 497 of 500 CSC-exposed spectra assigned correctly (Fig. 5D), and its largest weights fell at about 725, 1005 and 1042 cm^−1^, the same region as the strongest t-test bands. Spectra were vector-normalized, so differences reflect relative spectral contributions rather than absolute amounts.

**Figure 5.**
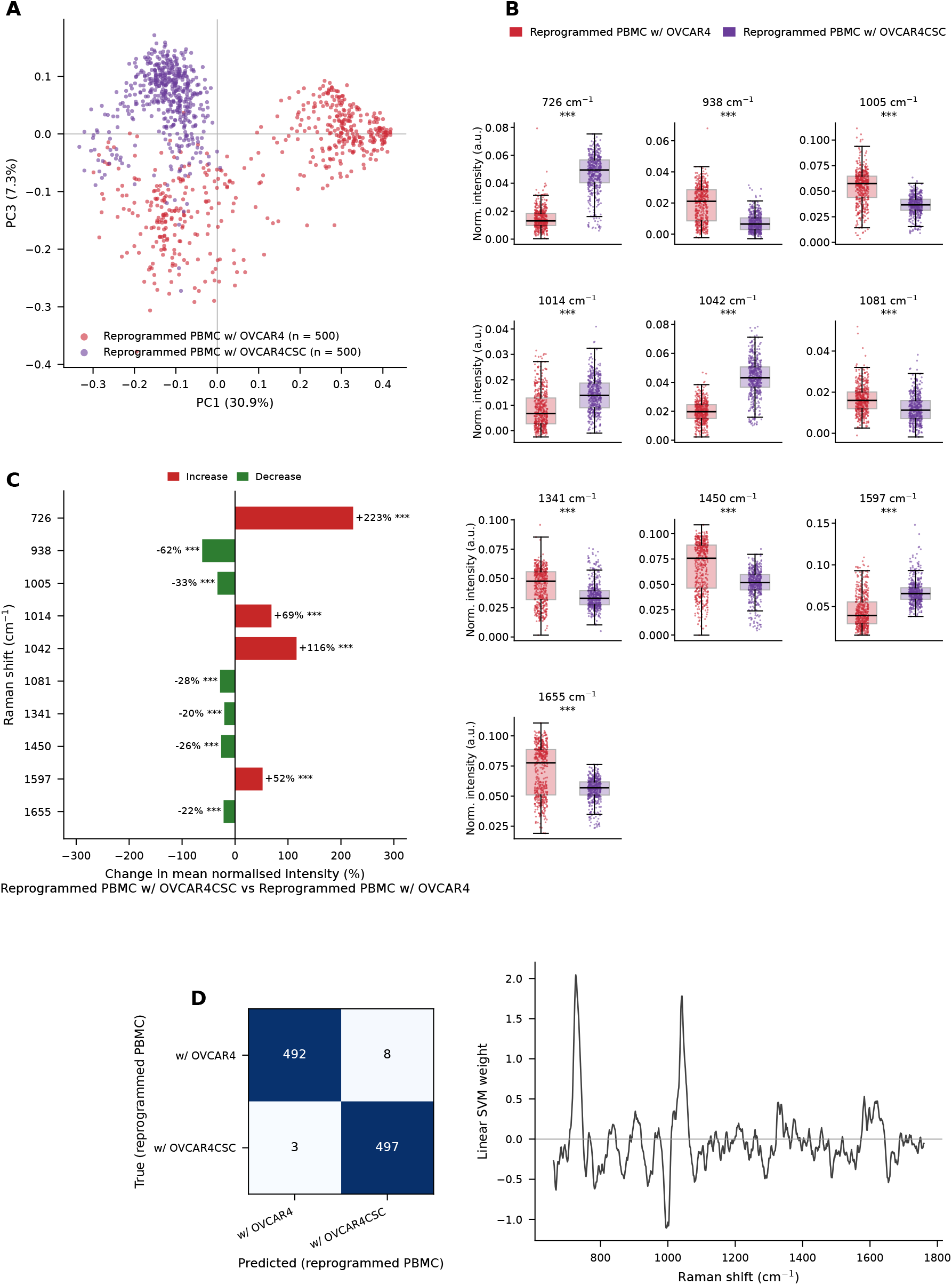
Reprogrammed PBMCs with OVCAR4 vs Reprogrammed PBMCs with OVCAR4-derived CSCs. (A) Principal component analysis (PCA). (B) t-test of the ten most significant bands. (C) Direction of change (% change in CSCs vs. reprogrammed PBMCs with OVCAR4-derived CSCs). (D) Support vector machine (SVM): confusion matrix and feature weights.

CSC-exposed PBMCs contributed more at 725 cm^−1^ (adenine ring breathing and phospholipid-associated modes, +223%), 1042 cm^−1^ (carbohydrate- and proline-associated region, +116%), 1014 cm^−1^ (phenylalanine-associated, +69%) and 1597 cm^−1^ (C=C and aromatic ring modes, +52%) (Fig. 5C)^14^. In contrast, OVCAR4-exposed PBMCs contributed more at 938 cm^−1^ (C– C backbone stretch of proline and valine, −62% in CSC-exposed), 1005 cm^−1^ (phenylalanine ring breathing, −33%), 1081 cm^−1^ (C–C/C–O of phospholipids and 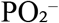 of nucleic acids, −28%), 1450 cm^−1^ (CH_2_/CH_3_ deformation of proteins and lipids, −26%), 1655 cm^−1^ (amide I, −22%) and 1341 cm^−1^ (guanine and CH deformation, −20%). The two conditions therefore differ in the balance between a phospholipid/carbohydrate-associated group of bands, which is higher after CSC exposure, and a protein-associated group, which is higher after OVCAR4 exposure. Given the overlap of several of these bands across biomolecule classes, we interpret them as shifts in overall composition rather than as evidence for a single molecule.

This pattern mirrors the difference between the cancer cells themselves: the bands higher in CSC-exposed PBMCs (725 and 1042 cm^−1^) correspond to those stronger in CSCs, and the protein-associated bands higher in OVCAR4-exposed PBMCs (1005, 1450 and 1655 cm^−1^) correspond to those stronger in OVCAR4 (Fig. 2), and both sets moved in the same directions when each condition was compared with untreated PBMCs (Figs. 3 and 4). This is consistent with each cancer cell type leaving a distinct, cell-type-related signature in PBMCs, for example through uptake of cancer-derived material, since ovarian cancer exosomes are known to alter lymphocyte signaling^10^, and with CSCs differing metabolically from the bulk tumor population^7,18,19^. It is equally consistent with residual cancer cells or debris in the PBMC preparation, which spectra alone cannot separate. Because Raman spectra of immune cells change with cell state^5–7^, the difference may also reflect distinct immune-cell responses, and lipid-associated increases of the type seen after CSC exposure have been linked to dysfunction in tumor-exposed immune cells^22,23^, although a single band cannot demonstrate lipid accumulation. The high classification accuracy shows that the groups are separable in this dataset; it does not show that the difference would generalize to independent experiments, since each condition was measured in one session. We note that these data do not identify the molecules or mechanism responsible.

## Conclusion

This study employed Surface-Enhanced Raman Scattering (SERS) on a nickel- and gold-coated substrate to compare ovarian cancer cells, their stem-like subset, and peripheral blood mononuclear cells (PBMCs) after co-culture with each. SERS demonstrated a clear distinction between OVCAR4 cells and OVCAR4-derived cancer stem cells (CSCs). OVCAR4 spectra exhibited a preference for protein-associated bands, including phenylalanine, amide III, CH_2_/CH_3_, and amide I. In contrast, CSC spectra exhibited a stronger presence of bands associated with phospholipid/nucleic acid and carbohydrate/glycogen regions^14,24^. After co-culture, PBMC spectra changed in both conditions, but in different ways. PBMCs co-cultured with OVCAR4 showed increased phenylalanine, amide I and CH_2_/CH_3_ bands, whereas PBMCs co-cultured with CSCs showed the largest increases in the phospholipid/nucleic-acid-associated 728 cm^−1^ band and the carbohydrate-associated 1045 cm^−1^ band. The two co-cultured PBMC populations were readily separable from each other (98.9% cross-validated SVM accuracy), and the bands that separated them corresponded to those that distinguished the two cancer cell types. These results show that exposure to each cancer cell type leaves a distinct, reproducible spectral signature in PBMCs and are consistent with transfer of cancer-cell-derived material^10^ or distinct immune-cell response^15,20,21^.

The spectra alone cannot identify the molecules involved or distinguish uptake of cancer-derived material from residual cancer cells or changes in cell composition within the mixed PBMC population. Additionally, each condition was measured in a single session, so the findings demonstrate separability within this dataset rather than reproducibility across independent experiments. SERS, however, provides a rapid, label-free method for detecting and comparing the effects of various cancer cell populations on immune cells.

